# Copy numbers of 45S and 5S ribosomal DNA arrays lack meaningful correlation in humans

**DOI:** 10.1101/2020.07.06.189753

**Authors:** Ashley N. Hall, Tychele N. Turner, Christine Queitsch

## Abstract

The ribosomal DNA genes are tandemly arrayed in most eukaryotes and exhibit vast copy number variation. There is growing interest in integrating this variation into genotype-phenotype associations. Here, we explored a possible association of rDNA copy number variation with autism spectrum disorder and found no difference between probands and unaffected siblings. However, rDNA copy number estimates from whole genome sequencing are error-prone, so we sought to use pulsed-field gel electrophoresis, a classic gold-standard method, to validate rDNA copy number genotypes. The electrophoresis approach is not readily applicable to the human 45S arrays due to their size and location on five separate chromosomes; however, it should accurately resolve copy numbers for the shorter 5S arrays that reside on a single chromosome. Previous studies reported tightly correlated, concerted copy number variation between the 45S and 5S arrays, which should enable the validation of 45S copy number estimates with CHEF-gel-verified 5S copy numbers. Here, we show that the previously reported strong concerted copy number variation is likely an artifact of variable data quality in the earlier published 1000 Genomes Project sequences. We failed to detect a meaningful correlation between 45S and 5S copy numbers in the large, high-coverage Simons Simplex Collection dataset as well as in the recent high-coverage 1000 Genomes Project sequences. Our findings illustrate the challenge of genotyping repetitive DNA regions accurately and call into question the accuracy of recently published studies of rDNA copy number variation in cancers and aging that relied on diverse publicly available resources for sequence data.

## Introduction

The genes encoding the ribosomal RNAs are present in long tandem arrays in most eukaryotes, often referred to as the ribosomal DNA. Most eukaryotic genomes contain two types of rDNA arrays: the 45S, encoding the 18S, 5.8S, and 28S rRNAs, and the 5S, encoding the 5S rRNA. Because of their repetitive nature, both rDNA arrays are susceptible to expansion and contraction, which can lead to vast copy number differences among individuals (1, 2). The phenotypic consequences of natural variation in rDNA copy number remain largely unexplored (3). An exception is the recent finding that, in humans, natural rDNA copy number variation is associated with altered global gene expression and mitochondrial abundance (1). Another study reported that the 45S and 5S arrays covary in copy number in mouse and human, an effect termed concerted copy number variation (4). Further, treating human cells with a drug causing nucleolar stress induced parallel loss of copies at both rDNA arrays. These findings were interpreted as evidence for natural selection maintaining concerted copy number variation, possibly in order to balance dosage of the 45S and 5S rRNAs.

Although dosage balance between the 45S and 5S rRNAs may facilitate efficient ribosome biogenesis, rRNA expression is not primarily a function of rDNA copy number (5). In many eukaryotes, a large proportion of the rDNA arrays is epigenetically silenced, forming the heterochromatic, dense core of the nucleolus. In exponentially growing human and yeast cells, about half of the rDNA copies are silenced (6, 7). In yeast, severe reduction of rDNA copy number results in all rDNA copies being expressed (5). Similarly, mutations interfering with epigenetic silencing cause the expression of additional rDNA copies (8, 9). Together, these data suggest that rRNA dosage is largely controlled through transcriptional regulation rather than by changing rDNA copy number, suggesting other selective pressures may lead to concerted copy number variation. If so, the reported strong concerted copy number variation would imply that the repetitive rDNA arrays undergo compensatory contractions or expansions across several distant genomic loci through yet undiscovered molecular mechanisms.

In addition to being an interesting biological phenomenon, concerted copy number variation holds promise for confirming the quality of rDNA copy number estimates. A key limit to incorporating rDNA copy number into genotype-phenotype associations is the ability to accurately quantify these genotypes. Because rDNA copy number estimates by short-read sequencing are error-prone due to batch effects (10), sequencing-based estimates should be validated by methods such as contour-clamped homogenous electric field gel electrophoresis (CHEF gels), which can separate megabase-sized DNA fragments. The human 45S arrays are too large and too numerous to be accurately quantified by CHEF gels, however the 5S can be readily measured (11). Combining 5S estimates by CHEF gels with a linear model relating 5S and 45S array copy numbers from sequencing data could permit the assessment of the accuracy of 45S rDNA copy number genotypes. Here, we sought to develop and use this method in order to facilitate the inclusion of rDNA copy numbers in genotype-phenotype association studies for autism spectrum disorder.

We analyzed concerted copy number variation between the 45S and 5S rDNA arrays in multiple datasets. These include the 163 low-coverage 1000 Genomes project samples in which concerted copy number variation was previously reported, as well as the far larger datasets of newly generated high quality, high-coverage sequencing data for both the 1000 Genomes Project and the Simons Simplex Collection. We estimated rDNA copy number by short read sequencing read depth in all three datasets. Unlike previously reported, we observe only weak correlations between the 45S and 5S rDNA arrays in the Simons Simplex Collection and high-coverage 1000 Genomes Project collection. We confirm that our analysis pipeline identifies the previously reported strong concerted copy number variation in the low-coverage 1000 Genomes Project data, and we show that concerted copy number variation is far weaker in the high-coverage data for the same samples. Furthermore, we show that copy number estimates between high and low-coverage data correlate poorly. The weak correlation between the 45S and 5S rDNA arrays is not an artifact of cell passaging between the initial 1000 Genomes Project sampling and the recent high-coverage resequencing effort because we observed the same result in the Simons Simplex Collection samples derived from blood. Both high-coverage datasets were generated at the New York Genome Center, using the same approaches to library preparation and sequencing. In contrast, the original, low-coverage 1000 Genomes Project data were generated in multiple genome centers using different methods. We recently reported that rDNA copy number estimation from whole-genome short-read sequencing data is sensitive to even subtle variation in sample processing and coverage in technical replicates (10). Our results on the lack of concerted copy number variation call into question several recently published associations of rDNA copy number with cancer and aging.

## Results

### Meaningful concerted copy number variation is not present in the Simons Simplex Collection

Our initial goal was to estimate rDNA copy number in the Simons Simplex Collection to determine if rDNA copy number is associated with autism spectrum disorder. Autism spectrum disorders associate with variants in hundreds of genes, including single nucleotide, tandem repeat, and copy number variants (12–14). rDNA copy number variation has not been assessed in autism spectrum disorder; however, it has been hypothesized that higher rDNA copy number associates with a more severe intellectual disability due to the increased potential for rRNA translation (15). The Simons Simplex Collection has sequenced hundreds of families with a child affected by an autism spectrum disorder (16–20). We estimated rDNA copy number in 7,268 individuals from families in which both of the parents, the proband, and an unaffected sibling were sequenced. We detected no difference in rDNA copy number based on autism status (**Figure 1A**), using read coverage estimates (4). We asked whether within probands, rDNA copy number associates with degree of intellectual disability by comparing individuals with an IQ of <70 to those with an IQ of >=70. We detected a statistically significant difference (nominal p=0.03) of individuals with an IQ<70 having on average eight more rDNA copies than those with IQ>=70, consistent with the published hypothesis. However, eight additional rDNA copies seem unlikely to be biologically relevant, given an average copy number of ~250. Moreover, we previously reported that rDNA copy number estimates based on short-read whole genome sequencing data can be error prone, with eight copies certainly being within the range of error (10). Therefore, we sought to validate rDNA copy number in a subset of samples by alternate methods.

**Figure 1:**
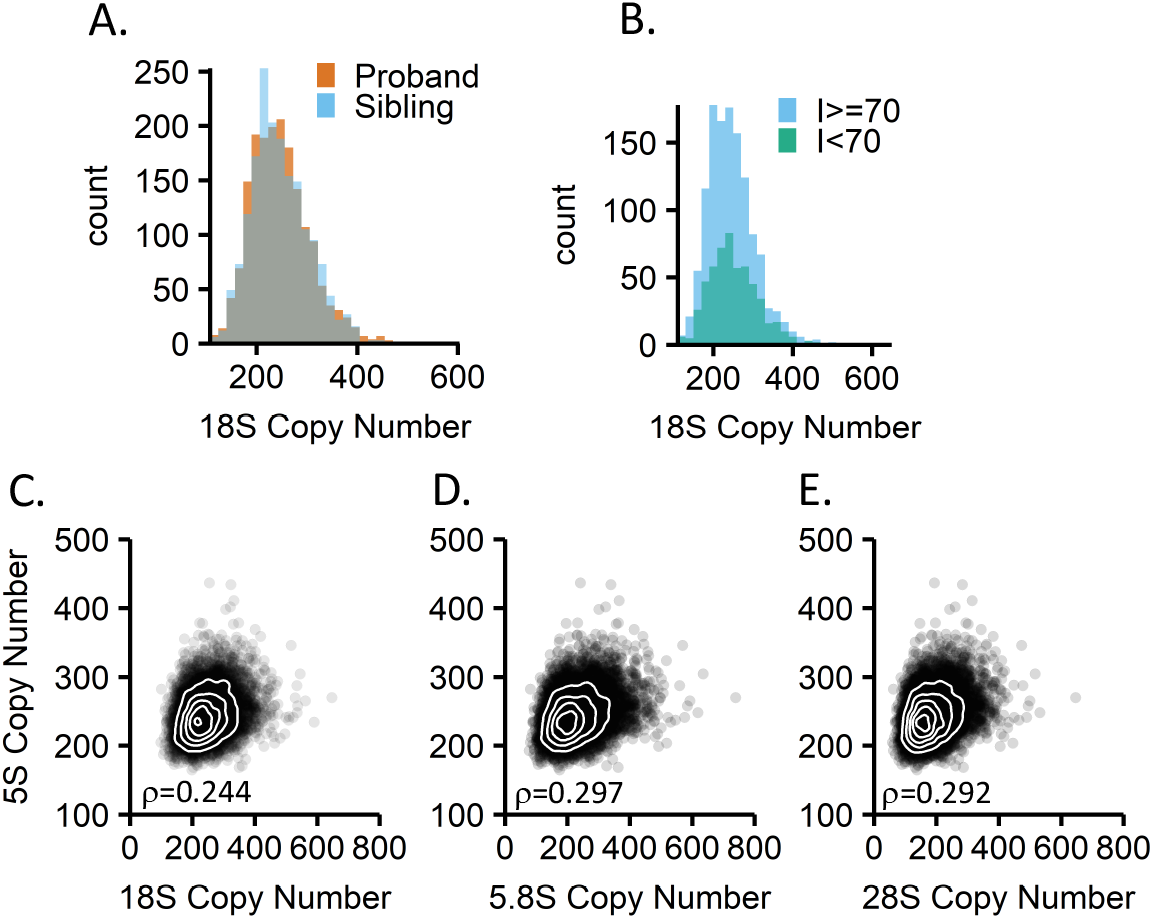
rDNA copy number estimates and correlations of 45S and 5S rDNA regions in the Simons Simplex Collection. A: rDNA copy number distributions for probands (n=1,774) and unaffected siblings (n=1,774) in the Simons Simplex Collection. Paired t-test p-value: 0.937. B: Comparison of rDNA copy number in probands with severely low IQ (I<70, n=548) or not (I>=70, n=1,244). C-E: Correlations of each the 18S, 5.8S, and28S regions of the 45S rDNA array to the 5S rDNA array with Spearman’s rho indicated (n=7,210).

Our preferred alternate method to estimate rDNA copy number is the CHEF gel. CHEF gels are the gold standard to estimate rDNA copy number, but they have limitations. In humans, the 45S rDNA array makes up the short arm of each of the 5 acrocentric chromosomes, so when assessing rDNA copy number by CHEF gel ten distinct bands should be observed. No previous study has ever observed ten distinct bands in a 45S CHEF gel, possibly because some rDNA loci are too large to be resolved (11). In contrast, the 5S rDNA, residing in a single locus, can be readily measured by CHEF gels. Given the previously reported concerted copy number variation between the 5S and 45S rDNA arrays (4), we planned to validate the 5S rDNA copy number estimates by CHEF gels to infer the accuracy of the sequencing-based 45S rDNA copy number estimates.

To this end, we first estimated 5S rDNA copy number in the Simons Simplex Collection to determine the strength of concerted copy number variation. We found weaker correlation than what was reported previously: The 18S and 5S copy numbers correlate with a Spearman coefficient of 0.24, while in the initial study using 1000 Genomes Project data the Spearman coefficient was 0.61 (**Figure 1C, Table 1**). As we were primarily interested in predicting 45S rDNA copy numbers from 5S rDNA copy numbers, we tested a linear model relating the copy numbers. This linear model is not predictive; the R^2^ value is 0.061. As expected, we observe similar trends when analyzing the 28S and 5.8S copy numbers in relation to the 5S number (**Figure 1 DE, Table 1**).

**Table 1:** Correlations between 45S and 5S rDNA copy numbers in the Simons Simplex Collection. The Spearman correlation coefficient and linear models that describe the relationships between the 45S and 5S rDNA copy numbers are shown.

| SSC (n=7210) |  |  |  |  |  |
| --- | --- | --- | --- | --- | --- |
| y | x | Spearman | Spearman p-value | Linear model | Multiple R-squared |
| 5S | 18S | 0.244 | < 2.2e-16 | $y=0.127x + 213$ | 0.061 |
| 5S | 5.8S | 0.297 | < 2.2e-16 | $y=0.134x + 214$ | 0.094 |
| 5S | 28S | 0.292 | < 2.2e-16 | $y=0.155x + 215$ | 0.093 |

### Concerted copy number variation in high-coverage 1000 Genomes data is weak

We next tested if weak concerted copy number variation was specific to the Simons Simplex Collection dataset. The 1000 Genomes Project recently released a new dataset of higher coverage sequencing data for approximately 2,500 samples (Michael Zody, personal communication). The sequencing data from the high-coverage dataset were generated by a single sequencing center with a single library preparation and sequencing method and a target genome coverage of ~30X. This sequencing center also generated the Simons Simplex Collection dataset. We estimated rDNA copy number in 2,419 of the high-coverage 1000 Genomes Project samples which displayed normal karyotypes. We observe a weak but significant correlation between each the 18S, 5.8S, and 28S copy numbers with the 5S copy number: The Spearman coefficients are 0.084, 0.111, and 0.118, respectively (**Fig.2A and Supplemental Figure 1, Table 2**). Despite the significance of the correlation, there is no predictive power to the relationship between the 45S and the 5S copy numbers: For example, a linear model relating the 18S and 5S copy numbers has an R^2^ value of 0.005 (**Table 2**).

**Figure 2:**
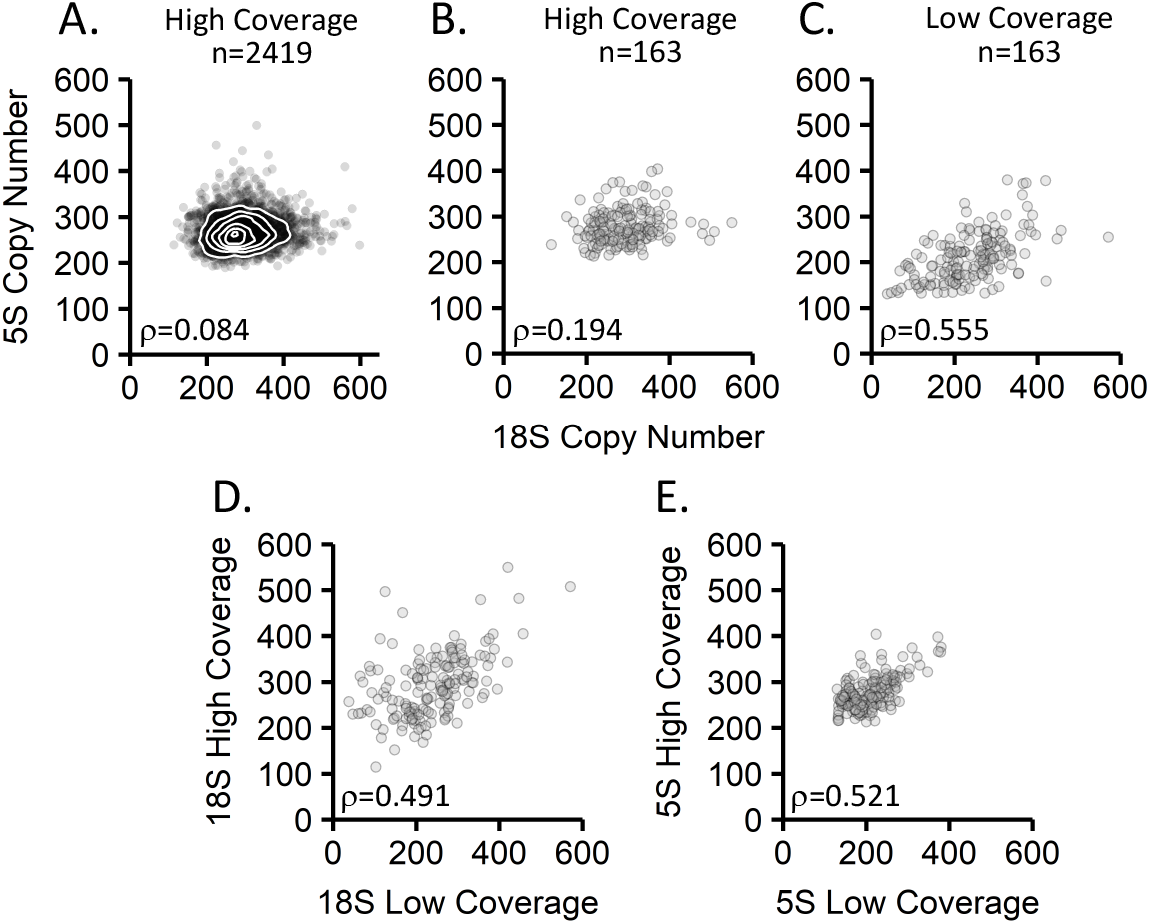
Correlations of the 5S and 45S rDNA copy numbers in 1000 Genomes Project data. A: Correlation of the 18S to the 5S in the high-coverage 1000 Genomes Project data (n=2,419). B: Correlation of the 18S to the 5S in the subset of 1000 Genomes Project data samples also analyzed in the low-coverage dataset (n=163). C: Correlation of the 18S to the 5S in the low-coverage 1000 Genomes Project data (n=163). D-E: Comparison of rDNA copy number estimates for the same cell lines sequenced separately in the high-coverage and low-coverage 1000 Genomes datasets for the 18S locus (D) and the 5S locus (E). Spearman’s rho is indicated in each panel, n=163.

**Table 2:** Correlations between 45S and 5S rDNA copy numbers in the high-coverage 1000 Genomes Project dataset. The Spearman correlation coefficient and linear models that describe the relationships between the 45S and 5S rDNA copy numbers are shown.

| High coverage data (n=2419) |  |  |  |  |  |
| --- | --- | --- | --- | --- | --- |
| y | x | Spearman | Spearman p-value | Linear model | Multiple R-squared |
| 5S | 18S | 0.084 | 3.69E-05 | $y = 0.037x + 258$ | 0.005 |
| 5S | 5.8S | 0.111 | 4.90E-08 | $y = 0.044x + 255$ | 0.010 |
| 5S | 28S | 0.118 | 6.97E-09 | $y = 0.061x + 255$ | 0.011 |

The weak concerted copy number variation signal in the high-coverage 1000 Genomes Project dataset is not simply due to an increased number of samples. If we exclusively analyze the high-coverage samples that were a part of the low-coverage study (n=163), we still observe a much weaker correlation than previously reported: the 18S and 5S correlate with a Spearman coefficient of 0.194 (**Figure 2B, Supplemental Figure 1, Table 3**). Our results raise the question whether differences in analysis methods or differences in data quality are responsible for the observed discrepancy with the previously reported findings (4).

**Table 3:** Correlations between 45S and 5S rDNA copy numbers in the subset of high-coverage 1000 Genomes Project data also analyzed in the low-coverage dataset. The Spearman correlation coefficient and linear models that describe the relationships between the 45S and 5S rDNA copy numbers are shown.

| High coverage data (n=163) |  |  |  |  |  |
| --- | --- | --- | --- | --- | --- |
| y | x | Spearman | Spearman p-value | Linear model | Multiple R-squared |
| 5S | 18S | 0.194 | 1.36E-02 | $y = 0.091x + 251$ | 0.029 |
| 5S | 5.8S | 0.217 | 5.73E-03 | $y = 0.091x + 251$ | 0.041 |
| 5S | 28S | 0.232 | 3.08E-03 | $y = 0.121x + 251$ | 0.042 |

### Our pipeline reproduces concerted copy number variation observed in low-coverage 1000 Genomes Project data

A key difference between our analysis and the original study in which concerted copy number variation was reported lies in the alignment pipelines and post-alignment corrections. We used BWA for alignment (21), while bowtie2 was used in the original study (4, 22). Additionally, the original study used a correction for pseudogene content, which we did not perform because no significant differences in concerted copy number variation were observed with or without corrections (4). To ensure that our analysis pipeline identifies the previously reported concerted copy number variation, we applied it to 163 of the 168 low-coverage 1000 Genomes Project samples previously studied.

Consistent with the original study, our analysis pipeline detected strong concerted copy number variation between the 45S and 5S loci in the low-coverage 1000 Genomes Project data. The 18S, 5.8S, and 28S rRNA gene copy numbers correlate to the 5S copy number with Spearman coefficients of 0.56, 0.79, and 0.69 (**Table 4, Figure 2C, Supplemental Figure 1**). These coefficients are similar to those previously published, which are 0.61, 0.80, and 0.73, respectively (4). We conclude that the failure to detect strong concerted copy number variation does not arise from differences in copy number estimation methods but is likely due to differences in the datasets used for analysis.

**Table 4:**
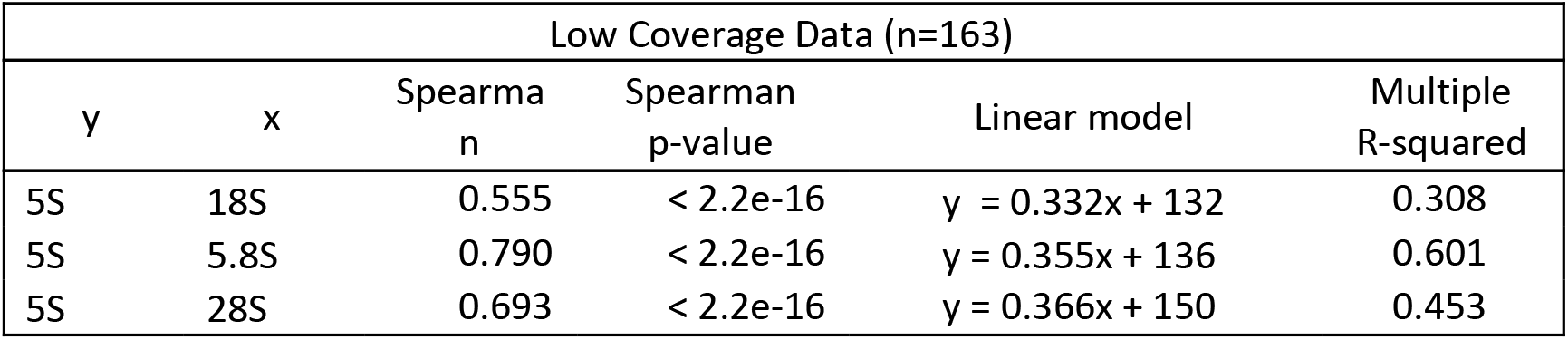
Correlations between 45S and 5S rDNA copy numbers in the low-coverage 1000 Genomes Project dataset. The Spearman correlation coefficient and linear models that describe the relationships between the 45S and 5S rDNA copy numbers are shown.

Indeed, we find that the low-coverage and high-coverage data yield different rDNA copy number estimates for the same cell lines. In comparing 18S estimates of the same cell lines in the two different datasets, the rDNA copy numbers only correlate with Spearman coefficients of 0.49. The 5S locus shows a Spearman correlation of 0.52 between the two datasets. These values are far lower than would be expected when analyzing the same cell lines (**Figure 2**). The scenario that rDNA copy numbers have changed between samplings for low- and high-coverage data generation is unlikely as there are several reports documenting that rDNA copy number is largely stable in cell lines (23, 24). Taken together, our results are consistent with previous reports that different library preparations of the same samples often yield different rDNA copy number estimates (10).

### Low-coverage 1000 Genomes Project sequencing data come from multiple sources

We wanted to further explore which differences in the low- and high-coverage datasets lead to different rDNA copy number estimates and the lack of meaningful concerted copy number variation. One difference between the datasets is that the low-coverage 1000 Genomes sequencing data were produced by any of seven sequencing centers, while the high-coverage data were produced by a single center. The seven sequencing centers used various library preparation methods (25), and library preparation methods and batch effects can influence rDNA copy number estimates (10). Moreover, some samples included reads from multiple library preparations and/or sequencing centers, turning the low-coverage copy number estimates into composite estimates from different libraries.

For each of the 163 low-coverage 1000 Genomes Project samples, we split the sequencing files by library preparation ID. Library depths varied by nearly two magnitudes: chromosome 1 coverage for individual libraries ranged from 0.24X to 14.4X, with an average of 5.4X. When we analyze rDNA copy number estimates from individual libraries by sequencing center, we find that some centers produced sequence data that biased rDNA copy estimates toward higher or lower values (**Supplemental Figure 2**). As it is unlikely that certain centers were assigned samples with abnormally high or low rDNA copy number, the observed bias likely arose through differences in library preparation methods. This bias was not observed in the high-coverage data for the same samples, which supports the notion that sample assignment was not biased.

Of the 163 low-coverage samples, rDNA copy number estimates for 54 samples were based on sequencing data generated from multiple libraries (**Figure 3**). Of these 54 samples, some samples, such as NA12154 and NA0700, contained multiple libraries that yielded 18S estimates very similar to both each other and our high-coverage data estimate. Others, such as NA11892 or NA12778, contained one library with an estimate very similar to our high-coverage data estimate but also other libraries with estimates near zero copies. The libraries with severe underestimates of rDNA copy number tended to have lower coverage, suggesting that read depth may influence rDNA copy number estimates.

**Figure 3:**
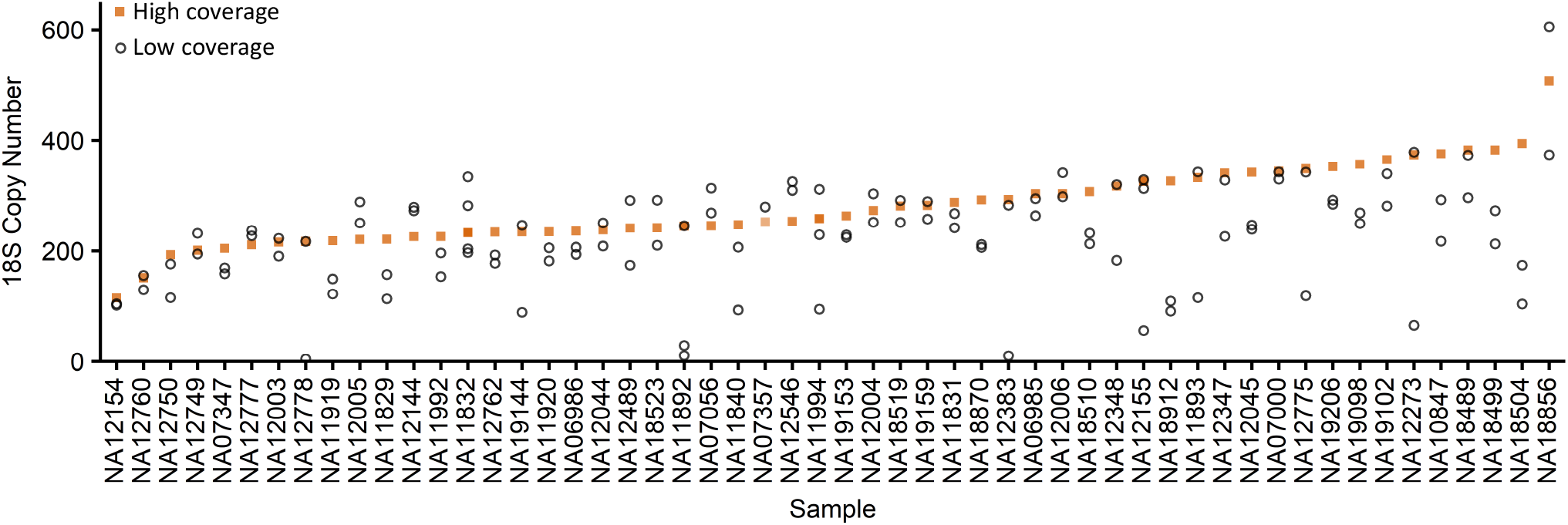
Comparison of 18S copy number estimates for different libraries made from the same cell lines. Of the low-coverage 1000 Genomes Project samples, 54 contained data from multiple sequencing libraries. Orange squares denote the high-coverage 1000 Genomes Project estimate. Open black circles indicate individual library estimates for each sample. Samples are ordered by high-coverage 18S estimate.

### Sequencing coverage does not account for the magnitude of rDNA copy number differences observed between the high and low-coverage 1000 Genomes Project datasets

We next investigated if depth of read coverage drives the differences in rDNA copy number estimation between the high- and low-coverage 1000 Genomes Project datasets. We performed downsampling experiments on four samples spanning the range of rDNA copy numbers estimates from the high-coverage 1000 Genomes Project dataset. Randomly downsampling of the high-coverage data to 400, 300, 200, 100, 50, and 10 million reads ten times each reveals that reducing coverage does affect rDNA copy number estimates to a small degree. Reassuringly, the average of ten independent downsamplings for a sample was close to the copy number estimate of the full dataset (**Figure 4**). For example, NA07357 had an estimated copy number of 252 in the high-coverage dataset. Its average copy number from downsampling ten times to 10 million reads was 249. Nevertheless, the range of copy number estimates increased with decreased coverage. At 10 million reads, the range of 18S estimates for NA07357 varied from 222 to 272, while at 100 million reads it varied from 244 to 260.

**Figure 4:**
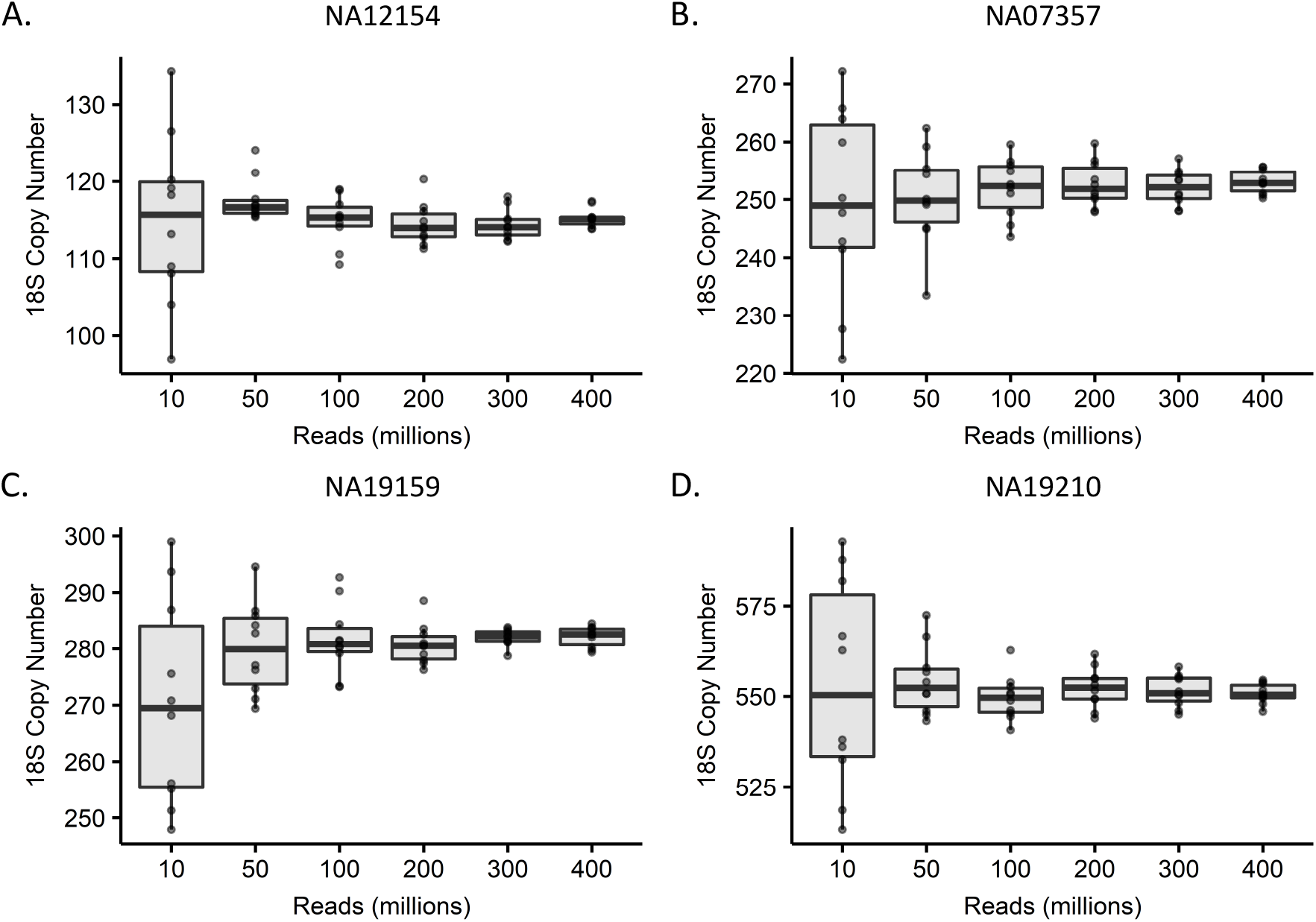
Read coverage downsampling of four high-coverage 1000 Genomes Project samples. A: NA12154 has an 18S copy number of 115 in the full dataset. B: NA07357 has an 18S copy number of 252 in the full dataset. C: has an 18S copy number of 282 in the full dataset. D: NA19210 has an 18S copy number of 550 in the full dataset. Note different Y axis scales for each graph. Ten independent downsamplings were performed per sample.

For comparison, the individual libraries for the low-coverage 1000 Genomes Project samples have a mean coverage of 5.4X for chromosome 1. The comparable samples in the downsampling experiment would be those with 100 million reads, which have a chromosome 1 coverage of approximately 4.5X. For the four samples analyzed, at ~4.5X coverage, the copy number estimates are at most +/− 13 copies of the full dataset estimate. Even at 10 million reads (~0.45X coverage), copy number estimates are at most +/− 43 copies of the full dataset estimate. Meanwhile, comparing the high- and low-coverage estimates, the low-coverage libraries underestimated the 18S by an average of 57 copies. The different libraries ranged from underestimating by 371 copies to overestimating by 110 copies as compared to the high-coverage data. These differences are much larger than what can be explained by differences in sequencing coverage alone. We conclude that the vast discrepancies in copy number estimates between datasets are not due to sequencing coverage, and hypothesize that the discrepancies may have arisen through batch effects or technical differences in library preparation.

### Simons Simplex Collection and high-coverage 1000 Genomes Project data are likely higher quality than the low-coverage 1000 Genomes Project data

To test the above hypothesis, we analyzed the correlation of copy number estimates of the three 45S components to each other, an approach previously used as a quality metric (1). Because these components are part of the same array, they should correlate highly. As expected, the high-coverage data showed somewhat higher intra-45S correlations than the low-coverage data. For example, the 18S and 28S copy numbers in the high-coverage dataset have a Spearman correlation of 0.97 (**Figure 5, Table 5**). In the 163 samples analyzed in both the high- and low-coverage data, the 18S and 28S copy numbers in the high-coverage dataset showed a Spearman correlation of 0.97 but has a Spearman correlation of 0.92 in the low-coverage dataset (**Figure 5, Tables 6 and 7**). This trend is reinforced by analysis of a linear model relating the two copy numbers. For the 163 samples analyzed in both 1000 Genomes Project datasets, the R^2^ value for the high-coverage dataset was 0.95 while it was 0.85 for the low-coverage dataset (**Tables 6 and 7**). The improved intra-45S correlations suggest that the rDNA copy number estimates in the high-coverage 1000 Genomes Project dataset are of higher quality.

**Figure 5:**
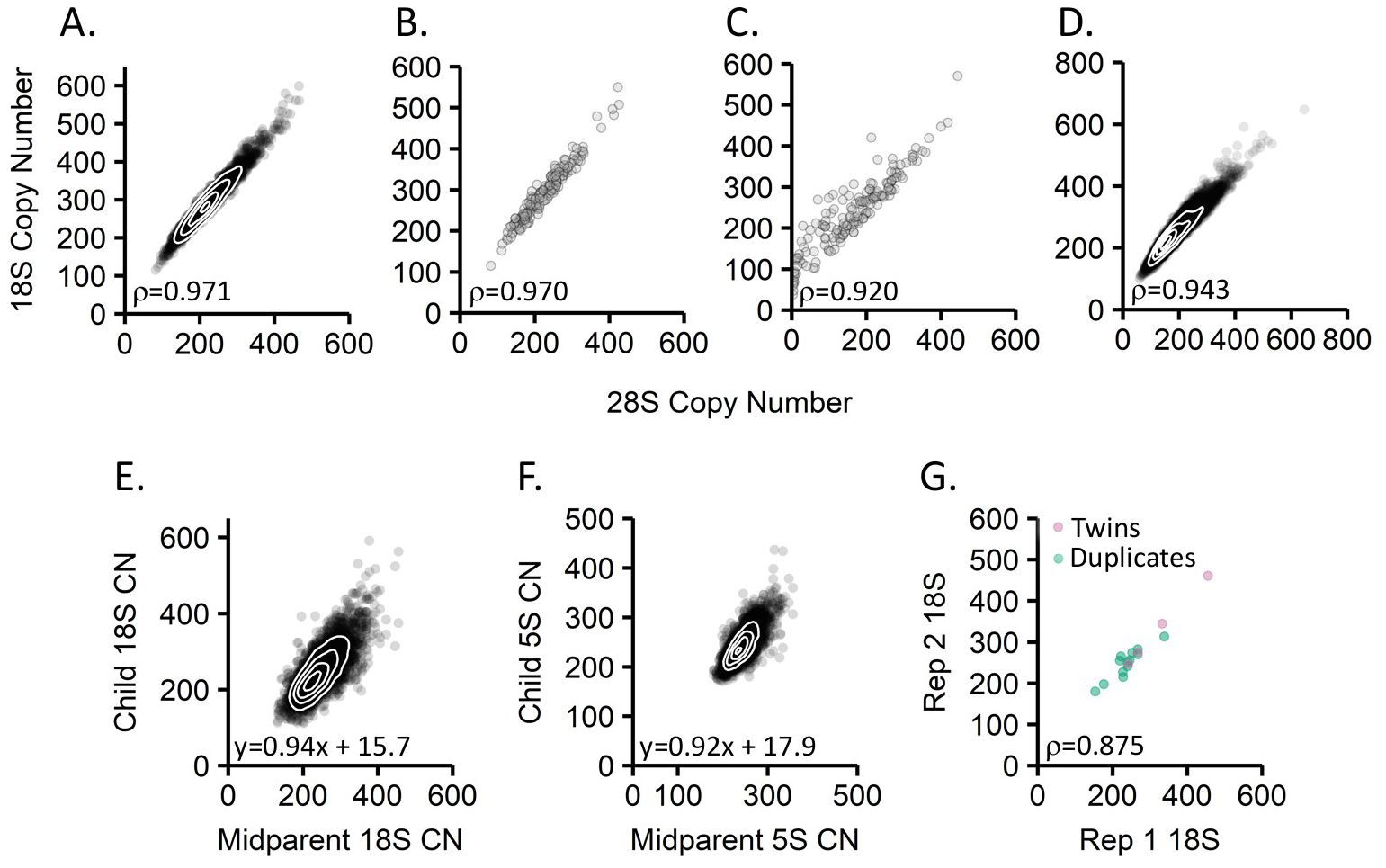
Data quality metrics for rDNA copy number estimates. A-D: Correlations between the 18S and 28S regions of the 45S rDNA repeat unit for the (A) high-coverage 1000 Genomes Project data (n=2,419), (B) subset of high-coverage 1000 Genomes Project data also analyzed in the low-coverage dataset (n=163), (C) low-coverage 1000 Genomes Project data (n=163), and (D) Simons Simplex Collection data (n=7,210). E: Heritability of the 18S copy number in the Simons Simplex Collection (n=3,548). F: Heritability of the 5S rDNA copy number in the Simons Simplex Collection. G: Comparison of 18S rDNA copy number estimates for either monozygotic twins (n=4 pairs) or for individuals sequenced twice in the Simons Simplex Collection (n=13). Spearman correlation indicated is for monozygotic twins and duplicates analyzed together.

**Table 5:**
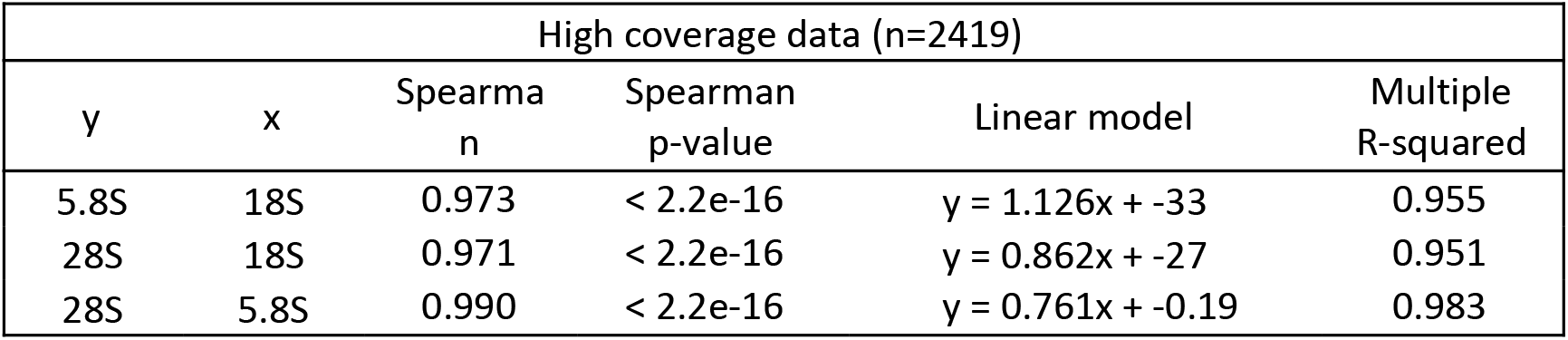
Correlations between the three rRNA genes encoded in the 45S repeat unit in the high-coverage 1000 Genomes Project dataset. The Spearman correlation coefficient and linear models that describe the relationships between the 18S, 5.8S, and 28S copy numbers are shown.

**Table 6:**
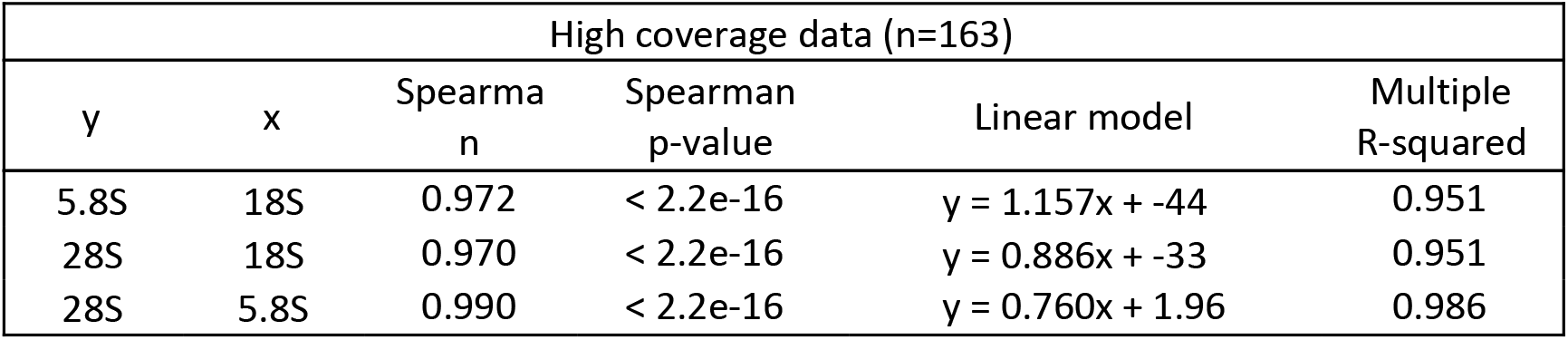
Correlations between the three rRNA genes encoded in the 45S repeat unit in the subset of high-coverage 1000 Genomes Project data also analyzed in the low-coverage dataset. The Spearman correlation coefficient and linear models that describe the relationships between the 18S, 5.8S, and 28S copy numbers are shown.

**Table 7:**
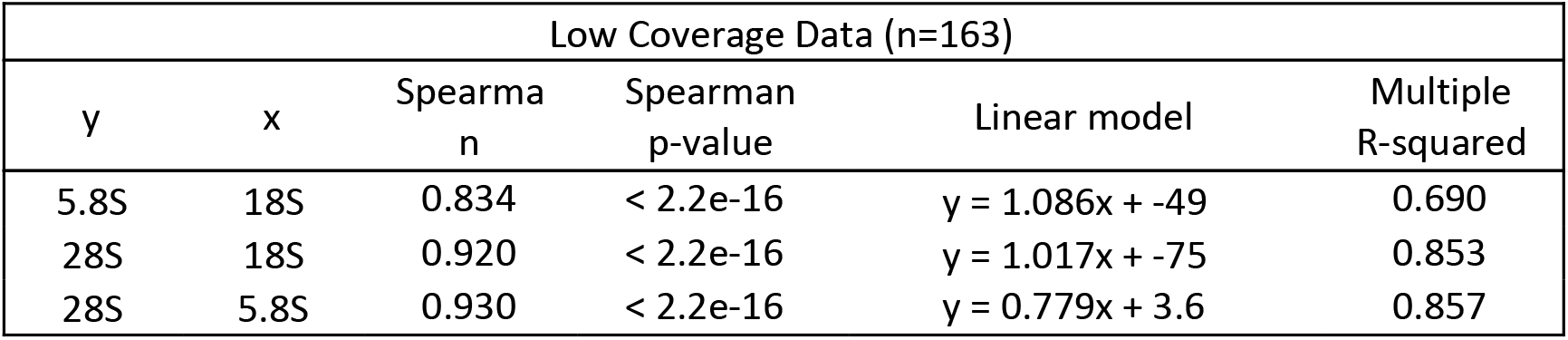
Correlations between the three rRNA genes encoded in the 45S repeat unit in the low-coverage 1000 Genomes Project dataset. The Spearman correlation coefficient and linear models that describe the relationships between the 18S, 5.8S, and 28S copy numbers are shown.

There are three different data quality metrics that can be analyzed with the Simons Simplex Collection data. As with the 1000 Genomes data, we first assessed intra-45S correlations, finding similarly high Spearman correlations: The 18S and 28S copy numbers showed a Spearman correlation of 0.943, and a linear model relating the two had an R^2^ value of 0.90. Unique to this study, we can also assess reproducibility of rDNA copy number estimates through use of monozygotic twins and duplicate samples. The Simons Simplex Collection includes four pairs of monozygotic twins that showed perfect correlation of copy number. Additionally, some individuals in the Simons Simplex Collection enrolled in other studies and were therefore sequenced twice, albeit by the same sequencing facility. These duplicate samples were identified from shared SNVs, and would be expected to share rDNA copy numbers. Although duplicate samples can show considerable deviation from one another due to technical issues (10), we found reasonably high correlation between the rDNA copy number estimates arising from the duplicated samples (Spearman correlation 0.81, **Figure 5F, Table 8**).

**Table 8:**
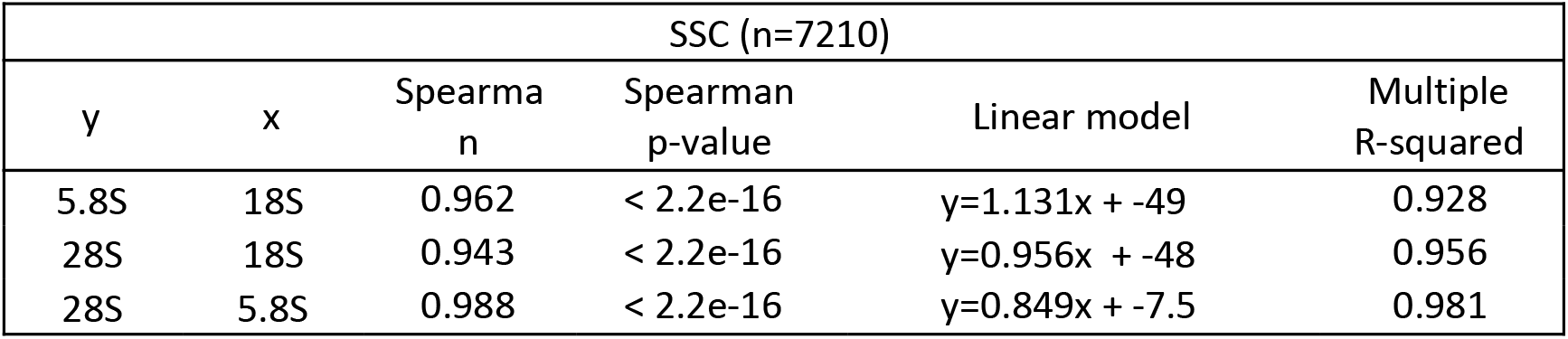
Correlations between the three rRNA genes encoded in the 45S repeat unit in the Simons Simplex Collection. The Spearman correlation coefficient and linear models that describe the relationships between the 18S, 5.8S, and 28S copy numbers are shown.

Finally, the family structure of the Simons Simplex Collection permits analysis of rDNA copy number heritability. We found high heritability of both 45S and 5S rDNA array components (h^2^=0.94 for the 18S, h^2^=0.92 for the 5S) (**Figure 5, Supplemental Figure 4**). Together, the heritability data, intra-45S correlations, and duplicated samples give confidence in the Simons Simplex Collection rDNA copy number estimates.

Taken together, we conclude that co-variation between the 45S and 5S rDNA arrays in both high-coverage datasets is weak and does not allow prediction of the copy number at one array based on the copy number at the other. This lack of meaningful concerted copy number variation holds true regardless of whether data were generated using lymphoblastoid cell lines or whole blood samples. The previously observed concerted copy number variation in the low-coverage dataset appears to be an artifact of lower data quality.

## Discussion

In this study, we sought to further explore the relationship between the copy numbers of the 45S and 5S rDNA arrays. We have previously reported that short-read sequencing estimates of rDNA copy number genotypes are error-prone (10). Given the previously published concerted copy number variation of the 5S and 45S rDNA arrays in humans, we thought this co-variation may provide a useful metric to predict 45S rDNA copy number from 5S copy number, the latter of which can be readily obtained by pulsed-field gel electrophoresis.

Working with two new, high-coverage sequencing datasets, we found weak concerted copy number variation that was not predictive. Sequencing coverage alone did not explain the discrepancy in copy number estimates or concerted copy number variation between samples sequenced in both the original low-coverage 1000 Genomes dataset and the newer high-coverage 1000 Genomes dataset. Available sequence data for many samples of the low-coverage dataset were derived from multiple library preparations from multiple sequencing centers. We suspect that differences in library preparation methods lead to these considerable discrepancies in rDNA copy number estimates.

In addition to the promise of a predictive model, the previously reported concerted copy number variation had fascinating implications for biology, in particular for genome maintenance and evolution. Selection for strongly concerted copy number variation suggests the existence of mechanisms that “count” and adjust copy numbers accordingly, operating across several genomic loci on separate chromosomes. A simpler explanation for keeping a specific ratio of 45S to 5S rDNA copies may be the much needed balance of 45S-encoded and 5S-encoded rRNA transcripts to build ribosomes efficiently. However, vast stretches of rDNA are typically silenced, forming the electron-dense heterochromatic nucleoli. In other words, the total number of rDNA copies does not necessarily reflect the number of transcribed copies. In yeast, the best characterized organism with regard to rDNA biology, more than half of the rDNA copies are silenced in wild type. In contrast, in mutant strains with few rDNA copies (~40), all copies are expressed to maintain wild-type ribosome biogenesis levels (5). The current consensus in this field is that rRNA dosage is not regulated by changes in copy number but rather by upregulated expression of euchromatic, transcriptionally active copies (5). In short, our finding that copy numbers at the 45S and 5S arrays show little correlation is biologically plausible. This lack of a meaningful correlation was previously observed in a large mutant collection of over 2000 strains and 40 natural isolates of *C. elegans*, using high-coverage sequence data to estimate rDNA copy number.

Some may argue that the observed differences in rDNA copy number between the low- and high-coverage 1000 Genomes datasets stem from biological differences; i.e. that the studied samples have acquired changes in rDNA copy number. We consider this an unlikely scenario. The DNA used for each sequencing effort was extracted from lymphoblastoid cell lines, which were propagated for an unknown number of generations from a common stock between each study. However, ribosomal DNA copy number has been reported as stable both in cell lines (23, 24) and multicellular organisms such as *C. elegans* (10, 26), except for when rDNA copy number is reduced to a level that causes fitness defects such as in *S. cerevisiae* or *D. melanogaster*.

We present our results as a cautionary tale about the challenges of genotyping repetitive DNA. While our results suggest that the high-coverage uniform sequencing performed by the New York Genome Center for the updated 1000 Genomes Project and the Simons Simplex Collection likely yielded more accurate rDNA copy number estimates, there is no certainty without validation through alternate methods. We are reassured by the fact that the observed lack of meaningful covariation between 45S and 5S rDNA copy numbers is consistent with current biological knowledge. We therefore trust our finding that there is no association of rDNA copy number with autism spectrum disorder. However, in light of our findings, we posit that care should be taken when drawing biological conclusions based on rDNA copy number estimates from short-read whole genome sequencing data. Changes in rDNA copy number have been reported for some cancers and in aging, prompting speculation about their role as drivers or essential players in both processes. Some of these findings may need to be re-evaluated by applying multiple data quality metrics to the analyzed sequence data, by conducting uniform, high-coverage re-sequencing, or by validating rDNA copy number through alternative approaches.

## Methods

### Alignment and copy number estimation

#### Sequence analysis

Samtools version 1.9 and bwa version 0.7.15 were used for all analyses.

Reference sequences for the 45S ribosomal DNA (U13369.1), 5S ribosomal DNA (X12811.1), mtDNA, and chromosome 1 of the human genome (GRCh38 reference) were downloaded from the NCBI nucleotide database with GenBank IDs as indicated in parenthesis. CRAM files were converted to fastq by Samtools fastq and aligned to the appropriate reference sequence by bwa mem with default parameters and converted to CRAM files with samtools view. Per-base read depth was calculated with Samtools depth outputting all positions, with the –d 0 flag to eliminate a maximum read depth cutoff.

To estimate ribosomal DNA copy number, average read depth across the whole 5S or 5.8S coding sequence was divided by the average chromosome 1 read depth. These regions correspond to positions 271-391 of X12811.1 for the 5S and positions 6623-6779 of U13369.1 for the 5.8S For the 18S and 28S subunits, segments of these genes previously used for concerted copy number variation analysis were used, and the average read depth at these regions was divided by the chromosome 1 depth (4). These are positions 3,841-3,985 of U13369.1 for the 18S, and 8,049-8,198 of U13369.1 for the 28S.

Copy number estimates, ribosomal DNA read coverage and average chromosome 1 coverage for all samples in this study are provided as supplemental tables 1-4.

#### Statistical analysis

Analyses were performed in Rstudio version 1.0.153, with R version 3.5.1.

## Supporting information

Supplemental Table 2: Low coverage 1000 Genomes rDNA copy numbers

Supplemental Table 1: High coverage 1000 Genomes rDNA copy numbers

Supplemental Table 3: SSC rDNA copy numbers

---

**Figure S1:**
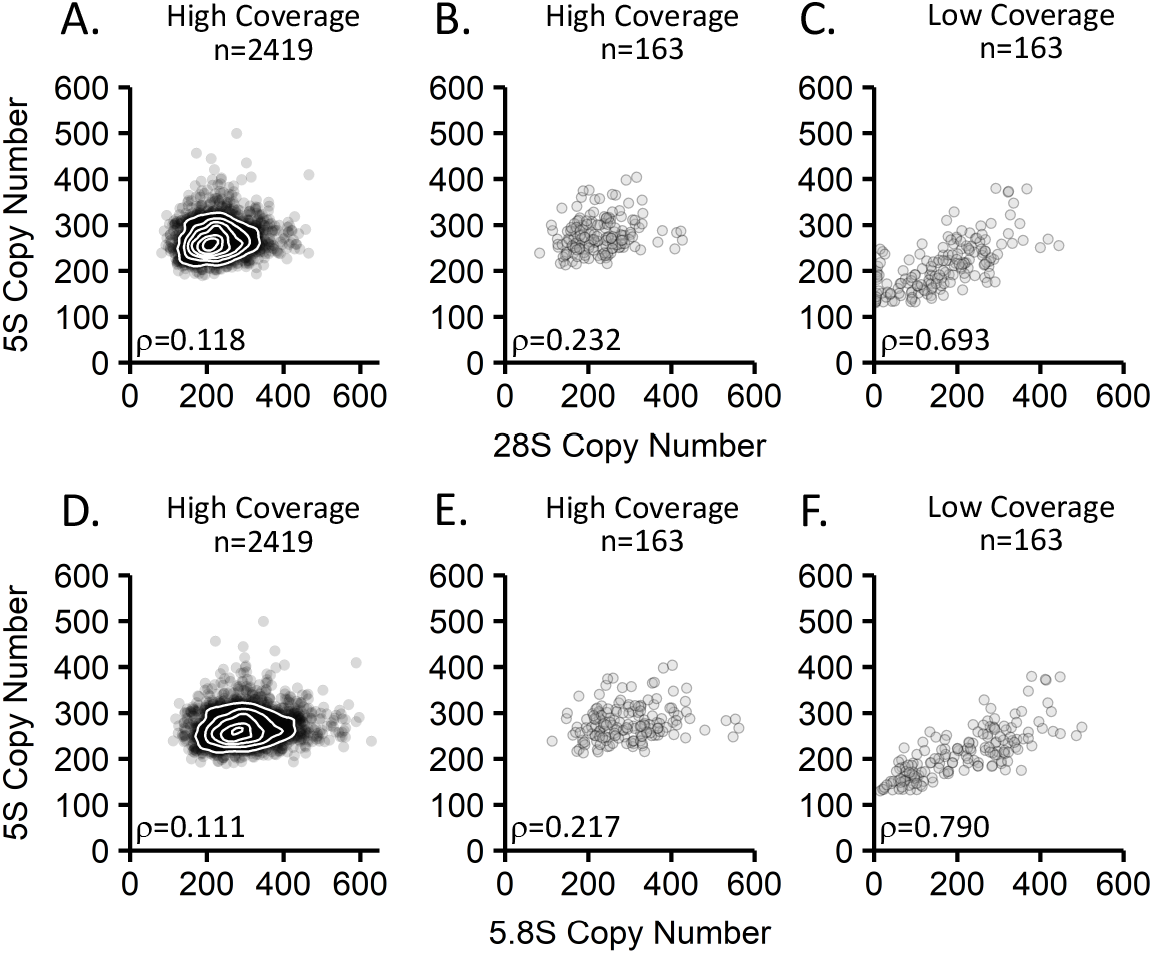
Correlations of rDNA copy numbers in 1000 Genomes Project data. A-C: Comparison of the 28S to 5S rDNA copy numbers in the (A) high-coverage 1000 Genomes Project data (n=2,419), (B) subset of high-coverage 1000 Genomes Project data also analyzed in the low-coverage dataset (n=163), and (C) low-coverage 1000 Genomes Project data (n=163). Bottom: Comparison of the 5.8S to the 5S rDNA copy numbers in the (D) high-coverage 1000 Genomes Project data (n=2,419), (E) subset of high-coverage 1000 Genomes Project data also analyzed in the low-coverage dataset (n=163), and (F) low-coverage 1000 Genomes Project data (n=163).

**Figure S2:**
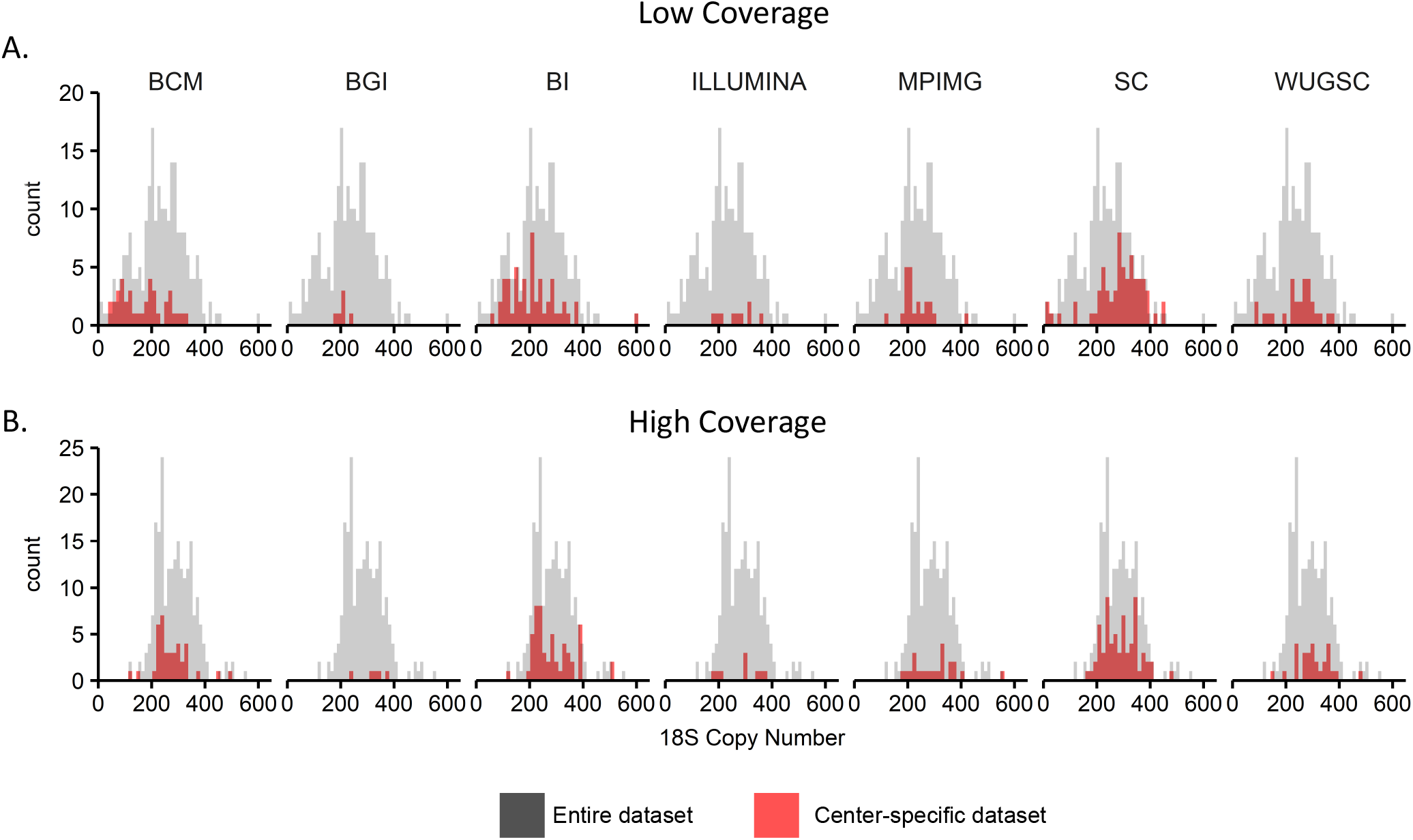
Distribution of 18S copy number estimates by sequencing center. A: Distribution of 18S copy number estimates from individual libraries sequenced by each of 7 sequencing centers (gray) for the 163 samples analyzed in the low coverage data (n=222 distinct libraries). Copy number estimates for samples produced by a given center in the low-coverage sequencing are highlighted in red. B: Distribution of 18S copy number estimates from the high-coverage 1000 Genomes Project data for the 163 samples analyzed in the low coverage data. The high-coverage data were not generated by any of these 7 centers: ‘B’ serves to demonstrate whether a center was assigned samples with a higher or lower copy number distribution. Samples that were composed of multiple distinct libraries in the low-coverage data are represented multiple times in the high-coverage plot.

**Figure S3:**
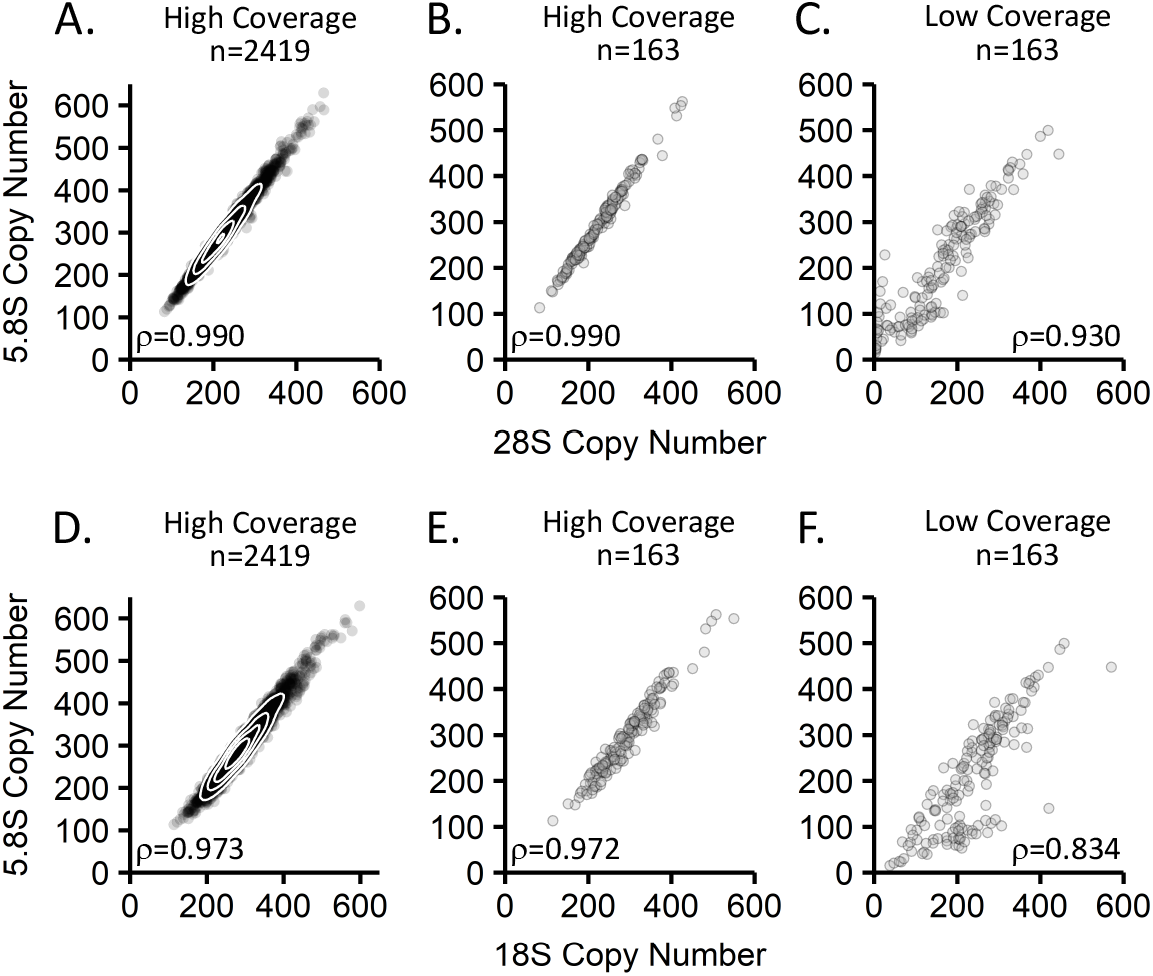
Comparison of copy number estimates of regions of the 45S rDNA repeat to each other. A-C: Comparison of the 28S to 5.8S rDNA copy numbers in the 1000 Genomes Project datasets. A: High-coverage 1000 Genomes Project data (n=2,419). B: Subset of high-coverage 1000 Genomes Project data also analyzed in the low-coverage dataset (n=163). C: Low-coverage 1000 Genomes Project data (n=163). D-F: Comparison of the 18S to the 5.8S rDNA copy numbers in the 1000 Genomes Project datasets. D: High-coverage 1000 Genomes Project data (n=2,419). E: Subset of high-coverage 1000 Genomes Project data also analyzed in the low-coverage dataset (n=163). F: Low-coverage 1000 Genomes Project data (n=163)

**Figure S4:**
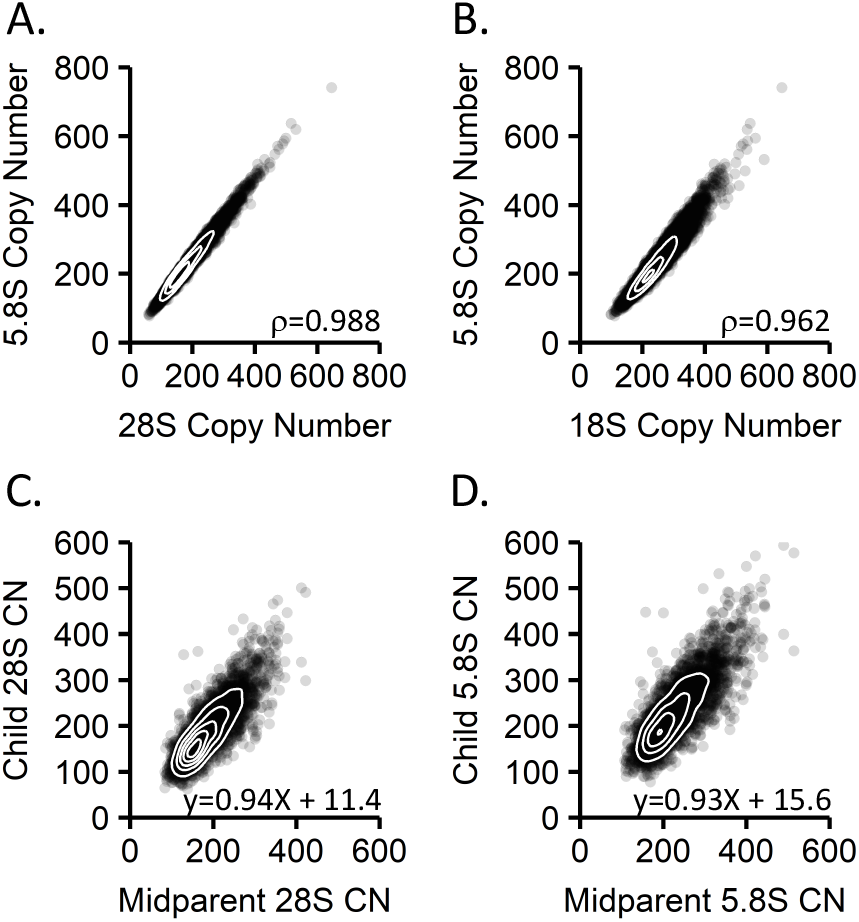
Data quality metrics for the Simons Simplex Collection. A: Comparison of the 28S to 5.8S rDNA copy numbers (n=7,210). B: Comparison of the 18S to 5.8S rDNA copy numbers (n=7,210). C: Heritability estimate of the 28S rDNA region (n=3,548). D: Heritability estimate of the 5.8S rDNA region (n=3,548).

## Data availability

High-coverage 1000 Genomes Project data are available at the following website: https://www.ebi.ac.uk/ena/data/view/PRJEB31736

The following cell lines/DNA samples were obtained from the NIGMS Human Genetic Cell Repository at the Coriell Institute for Medical Research: [NA06984, NA06985, NA06986, NA06989, NA06994, NA07000, NA07037, NA07048, NA07051, NA07056, NA07347, NA07357, NA10847, NA10851, NA11829, NA11830, NA11831, NA11832, NA11840, NA11843, NA11881, NA11892, NA11893, NA11894, NA11918, NA11919, NA11920, NA11930, NA11931, NA11932, NA11933, NA11992, NA11994, NA11995, NA12003, NA12004, NA12005, NA12006, NA12043, NA12044, NA12045, NA12046, NA12058, NA12144, NA12154, NA12155, NA12156, NA12234, NA12249, NA12272, NA12273, NA12275, NA12282, NA12283, NA12286, NA12287, NA12340, NA12341, NA12342, NA12347, NA12348, NA12383, NA12399, NA12400, NA12413, NA12414, NA12489, NA12546, NA12716, NA12717, NA12718, NA12748, NA12749, NA12750, NA12751, NA12760, NA12761, NA12762, NA12763, NA12775, NA12776, NA12777, NA12778, NA12812, NA12813, NA12814, NA12815, NA12827, NA12828, NA12829, NA12830, NA12842, NA12843, NA12872, NA12873, NA12874, NA12878, NA12889, NA12890]. These data were generated at the New York Genome Center with funds provided by NHGRI Grant 3UM1HG008901-03S1.

Sequencing data for the SSC is available through the Simons Foundation for Autism Research Initiative (SFARI) and is available to approved researchers at SFARI base (http://base.sfari.org, accession IDs: SFARI_SSC_WGS_p, SFARI_SSC_WGS_1, and SFARI_SSC_WGS_2).

## Acknowledgements

This work was supported, in part, by grants from the US National Institutes of Health (R00 MH117165 to T.N.T., F31 AG063450 to A.N.H, NIGMS grant R01 GM122088 to C.Q., and NHGRI grant 1RM1HG010461 to C.Q)

We are grateful to all of the families at the participating Simons Simplex Collection (SSC) sites, as well as the principal investigators (A. Beaudet, R. Bernier, J. Constantino, E. Cook, E. Fombonne, D. Geschwind, R. Goin-Kochel, E. Hanson, D. Grice, A. Klin, D. Ledbetter, C. Lord, C. Martin, D. Martin, R. Maxim, J. Miles, O. Ousley, K. Pelphrey, B. Peterson, J. Piggot, C. Saulnier, M. State, W. Stone, J. Sutcliffe, C. Walsh, Z. Warren, E. Wijsman). We appreciate obtaining access to genetic data on SFARI Base. Approved researchers can obtain the SSC population dataset described in this study SFARI_SSC_WGS_p, SFARI_SSC_WGS_1, and SFARI_SSC_WGS_2 by applying at https://base.sfari.org.

